# Massive variation of short tandem repeats with functional consequences across strains of *Arabidopsis thaliana*

**DOI:** 10.1101/145128

**Authors:** Maximilian O. Press, Rajiv C. McCoy, Ashley N. Hall, Joshua M. Akey, Christine Queitsch

## Abstract

Short tandem repeat (STR) mutations may be responsible for more than half of the mutations in eukaryotic coding DNA, yet STR variation is rarely examined as a contributor to complex traits. We assess the scope of this contribution across a collection of 96 strains of *Arabidopsis thaliana* by massively parallel STR genotyping. We found that 95% of examined STRs are polymorphic, with a median of six alleles per STR in these strains. Modest STR expansions are found in most strains, some of which have evident functional effects. For instance, three of six intronic STR expansions are associated with intron retention. Coding STRs are depleted of variation relative to non-coding STRs, consistent with the action of purifying selection, and some STRs show hypervariable patterns consistent with diversifying selection. Finally, we detect dozens of novel STR-phenotype associations that could not be detected with SNPs alone, validating several with follow-up experiments. Our results demonstrate that STRs comprise a large, unascertained reservoir of functionally relevant genomic variation.

## Introduction

Mean rates of mutation vary within genomes by several orders of magnitude (1), from ~10^−8^-10^−9^ for substitutions to 10^−3^−10^−4^ for short tandem repeat (STR) mutations (2–4). STR mutations generally occur through the addition or subtraction of repeat units. Given the prevalence of STR loci in eukaryotic genomes, we would expect more *de novo* STR mutations than single nucleotide substitutions in the human genome per generation (4), even in exonic regions (Table 1). Thus, while the overall mutation rate is under the strong control of natural selection, some loci experience many more mutations than others (5). The existence of such highly mutable loci violates simplifying assumptions of the infinite sites model of population genetics (6), namely that no locus mutates more than once in a population, as well as quantitative genetic models assuming contributions from many independent loci (7).

**Table 1.** Expected number of human mutations of various classes. Some plausible scenarios are bolded.

| Substitution rate<sup>a</sup> | STR mutation rate<sup>b</sup> | Exome size (bp)<sup>c</sup> | Exonic STRs<sup>d</sup> | Substitutions | STR mutations | log10(STR mutations / substitutions) |
| --- | --- | --- | --- | --- | --- | --- |
| $1.2 * 10^{-8}$ | $10^{-5}$ | $3 * 10^8$ | 5000 | 0.36 | 0.05 | -0.86 |
| **$1.2 * 10^{-8}$** | **$10^{-4}$** | **$3 * 10^8$** | **5000** | **0.36** | **0.5** | **0.14** |
| $1.2 * 10^{-8}$ | $10^{-3}$ | $3 * 10^8$ | 5000 | 0.36 | 5 | 1.14 |
| $1.2 * 10^{-8}$ | $10^{-5}$ | $3 * 10^8$ | 2000 | 0.36 | 0.02 | -1.26 |
| $1.2 * 10^{-8}$ | $10^{-4}$ | $3 * 10^8$ | 2000 | 0.36 | 0.2 | -0.26 |
| **$1.2 * 10^{-8}$** | **$10^{-3}$** | **$3 * 10^8$** | **2000** | **0.36** | **2** | **0.74** |
| $1.2 * 10^{-8}$ | $10^{-5}$ | $3 * 10^8$ | 1000 | 0.36 | 0.01 | -1.56 |
| $1.2 * 10^{-8}$ | $10^{-4}$ | $3 * 10^8$ | 1000 | 0.36 | 0.1 | -0.56 |
| **$1.2 * 10^{-8}$** | **$10^{-3}$** | **$3 * 10^8$** | **1000** | **0.36** | **1** | **0.44** |

**a:** Rate taken from refs. (8, 9).
**b:** Range of rates taken from refs. (2, 10). Different rates represent different points in the range. This value will vary according to the definition employed to select the set of STRs.
**c:** Approximate exome size taken from ref. (11).
**d:** According to ref. (12) there are 59,050 exonic STRs of at least three units, a very lax threshold. Values shown represent different possible subsets of exonic STRs showing non-trivial mutation rates.

In spite of the large effects that STRs can have on complex traits and diseases in model organisms and humans (13–15), their variation is rarely considered in genotype-phenotype association studies due to technical obstacles. However, STR genotyping methods of sufficient accuracy, throughput, and cost-effectiveness to ascertain STR alleles at high throughput have recently become available (16–18). Studies leveraging these methods suggest considerable contributions of STRs to heritable phenotypic variation (16, 19).

From an evolutionary perspective, the high mutation rate and clear phenotypic effects of STRs have been speculated to provide readily accessible evolutionary paths for rapid adaptation (20–22). On longer time scales, the presence of highly mutable STRs within coding regions appears to be maintained by selection (23–25). These observations independently argue for important roles of STRs as a reservoir of functional genetic variation.

In the present study, we apply massively parallel STR genotyping to a diverse panel of well-characterized *A. thaliana* strains. We use these data to generate and test hypotheses about the functional effects of STR variation, combining observations of gene disruption by STR expansion, inferences about STR conservation, and analyses of phenotypic association. Based on our results, we argue that STRs must be included in any comprehensive account of phenotypically relevant genomic variation.

## Results

### STR genotyping reveals complex allele frequency spectra

We targeted 2,050 STR loci for genotyping with molecular inversion probes (MIPs) (16) across a core collection of 96 *A. thaliana* strains (Methods). These loci were all less than 200 bp in length and had nucleotide purity of at least 90%, encompassing nearly all gene-associated STRs and ~40% of intergenic STRs (Fig. 1a). We used comparisons with the Col-0 reference genome, PCR analysis of selected STRs, and dideoxy sequencing to estimate that MIP STR genotype calls were ~95% accurate, and inaccurate calls were generally only one to two units away from the correct copy number (Supplementary Note, Supplementary Fig. 1-4, Supplementary Table 1).

**Figure 1.**
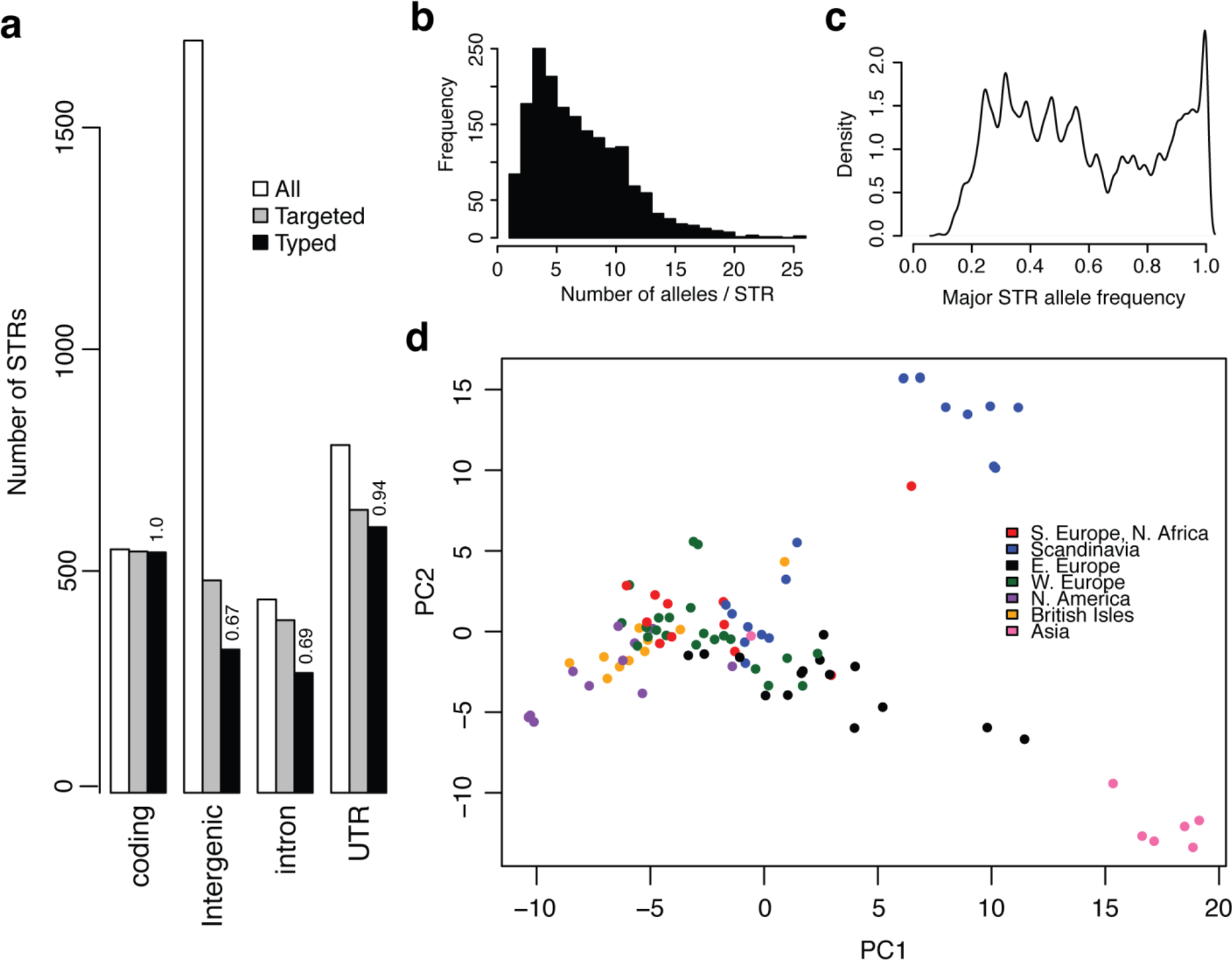
STRs in *A. thaliana* show a complex allele frequency distribution and geographic differentiation. (a): Distribution and ascertainment of STR loci. “All”: all STRs matching the definition of STRs for this study, *e.g.* <=180 bp length in TAIR10, >=89% purity in TAIR10, 2-10 bp nucleotide motif. “Targeted”: the 2050 STRs targeted for MIP capture. “Typed”: STRs successfully genotyped in the Col-0 genome in a MIPSTR assay. Numbers above bars indicate the proportion of targeted STRs in the relevant category that were successfully genotyped. (b): The distribution of allele counts across all genotyped STRs. (c): The distribution of major allele frequencies (frequency of the most frequent allele at each locus) across genotyped STRs. (d): Principal component analysis (PCA) reveals substantial geographic structure according to STR variation. PC1 and PC2 correspond, respectively, to 5.2% and 4.0% of total STR allele variance.

Across genotyped loci, we observed that 95% of STRs were polymorphic. Most STRs were highly multiallelic (Fig. 1b; mean=6.4 alleles, median=6 alleles), and this variation was mostly unascertained by the 1001 Genomes resource for *A. thaliana* (Supplementary Fig. 3a). Coding STRs were only slightly less polymorphic (mean=4.5 alleles, median=4 alleles) than non-coding STRs, though it is unknown whether this difference is due to purifying selection or variation in mutation rates. Highlighting the massive variation segregating at STR loci, 45% of STRs had a major allele with frequency < 0.5. This complicates the familiar concepts of major and minor alleles, which have provided a common framework for detecting genotype associations (Fig. 1c). Specifically, the Col-0 reference strain carries the major STR allele at only 48% of STR loci. Moreover, rarefaction analysis implied that more STR alleles at these loci are expected with further sampling of *A. thaliana* strains (Supplementary Fig. 4c).

Principal component analysis of STR variation revealed genetic structure corresponding to Eurasian geography (Fig. 1d, Supplementary Fig. 5), consistent with previous observations that genetic population structure is correlated with the geographic distribution of *A. thaliana* (26, 27). By corroborating previous observations from a much larger set of genome-wide single nucleotide polymorphism markers, this result demonstrates that a comparatively small panel of STRs suffices to capture detailed population structure in *A. thaliana*.

### Novel STR expansions are associated with splice disruptions.

We next examined the frequency and functional consequences of STR expansions in *A. thaliana.* STR expansions are high-copy-number variants of comparatively short STRs that are widely recognized as contributors to human diseases (28) and other phenotypes (29). While large (>150 bp) expansions are difficult to infer, we detected modest STR expansions using a simple heuristic that compares the longest allele to the median allele observed at each locus (Fig. 2a). We identified expansions in 64 of 96 *A. thaliana* strains, each carrying at least one expanded STR allele from one of 28 expansion-prone STRs (9 coding, 6 intronic, 8 UTR, 5 intergenic). Most expansions were found in multiple strains (Fig. 2b), although expansion frequency was likely underestimated due to a higher rate of missing data at these loci. STR expansions causing human disease can be as small as 25 copies (28), whereas we detected expansions up to ~50 copies.

**Figure 2.**
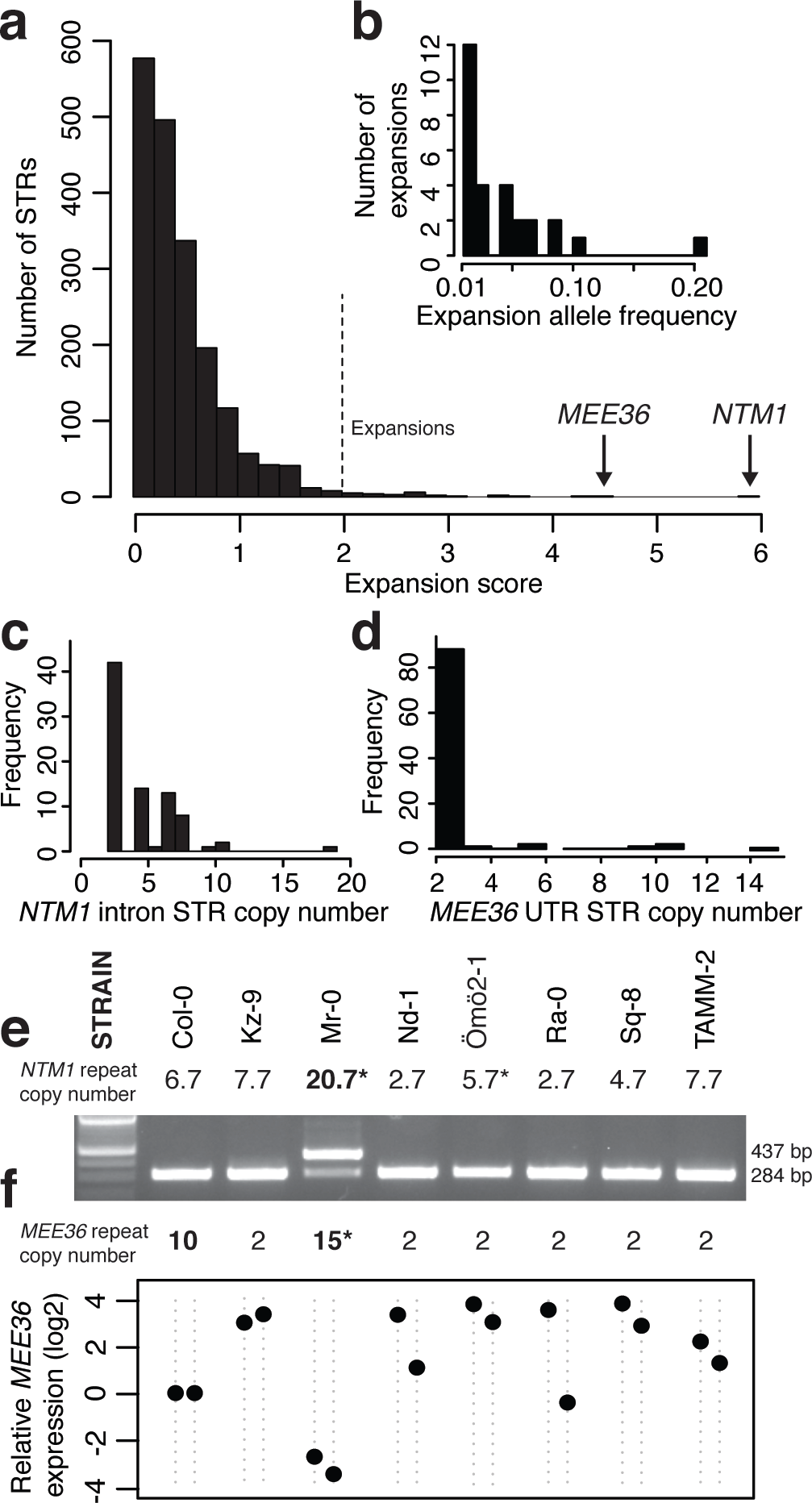
Inferring and assessing the functional effects of modest STR expansions. (a): The distribution of expansion scores across STRs, where the expansion score is computed as (max(STR length) − median(STR length)) / median STR length. We called any STRs with a score greater than 2 a modest expansion (indicated). (b): Distribution of allele frequencies of the 28 expanded STR alleles. (c, d): Distribution of STR copy number of the intronic STR in the *NTM1* gene and the 3’ UTR STR in the *MEE36* gene. (e): RT-PCR demonstrates intron retention in *NTM1* mRNA in the Mr-0 strain, which carries the STR expansion, yielding an aberrant 437-bp product. (f): *MEE36* transcript abundances measured by qRT-PCR and normalized relative to *UBC21* transcript levels. For each strain, two independent biological replicates are shown as points. Transcript levels are expressed relative to Col-0 levels (set to 1). *: STR genotype corrected by dideoxy sequencing. Strains and order are the same between (e) and (f).

We assayed the effects of STR expansions on expression of associated genes. The most dramatic expansions (with large relative copy number increase) affected an intronic STR in the *NTM1* gene (Fig. 2c; five other expansions also resided in introns) and a STR in the 3’ UTR of the *MEE36* gene (Fig. 2d). These genes, respectively, have roles in cell proliferation and embryonic development. Intronic STR mutations can disrupt splicing, causing altered gene function (30) and human disease (31). Therefore, we assayed effects on splicing of all six expanded intronic STRs. In three cases, the expanded allele was associated with partial or full retention of its intron (Supplementary Fig. 6, Supplementary Note), including the major *NTM1* splice form in the Mr-0 strain (Fig. 2e). We confirmed intron retentions by dideoxy sequencing of cDNA (Supplementary Fig. 7a). Retention of the *NTM1* intron is predicted to lead to a nonsense mutation truncating most of the NTM1 protein (Supplementary Fig. 7b). For the other two retentions, more complex and STR allele-specific mRNA species were formed (Supplementary Fig. 6, Supplemental Text). The *MEE36* STR expansion alleles were associated with dramatically reduced *MEE36* transcript levels (Fig. 2f), possibly due to the STR expansion altering transcript processing (32). These examples emphasize the potential for previously unascertained STR variation to modify gene function. Moreover, we show that the distribution of allele sizes itself can be informative, enabling predictions about STR functional effects.

### Signatures of functional constraint on STR variation.

Using the observed STR allele frequency distributions, we next attempted to infer selective processes acting on STRs. While previous models for evaluating functional constraint on STRs are few (33), there is consensus that selection shapes STR variation (21, 23, 33, 34). Naively, we would expect that coding STRs should show increased constraint (lower variation). Consistent with this expectation, we observed that most invariant STRs are coding (53 of 84 invariant STRs genotyped across at least 70 strains; odds ratio = 4.5, p = 5*10^−11^, Fisher’s Exact Test). However, methods of inferring selection by allele counting are confounded by population structure and mutation rate, which vary widely across STRs in this (Fig. 3a) and other studies (3, 35).

Therefore, to account for mutation rate and population structure, we used support vector regression (SVR) to predict STR variability across these 96 strains, using well-established correlates of STR variability (*e.g.* STR unit number and STR purity; Online Methods) (3, 10, 36). Selection was defined as deviation from expected variation of a neutral STR among strains. We trained SVRs on the set of intergenic STRs, which should experience minimal selection relative to STRs associated with genes (Supplementary Fig. 8-10). We used bootstrap aggregation of SVR models to compute a putative constraint score for each STR by comparing its observed variability to the expected distribution from bootstrapped SVR models (Fig. 3b, Supplementary Note, Supplementary File 1).

**Figure 3.**
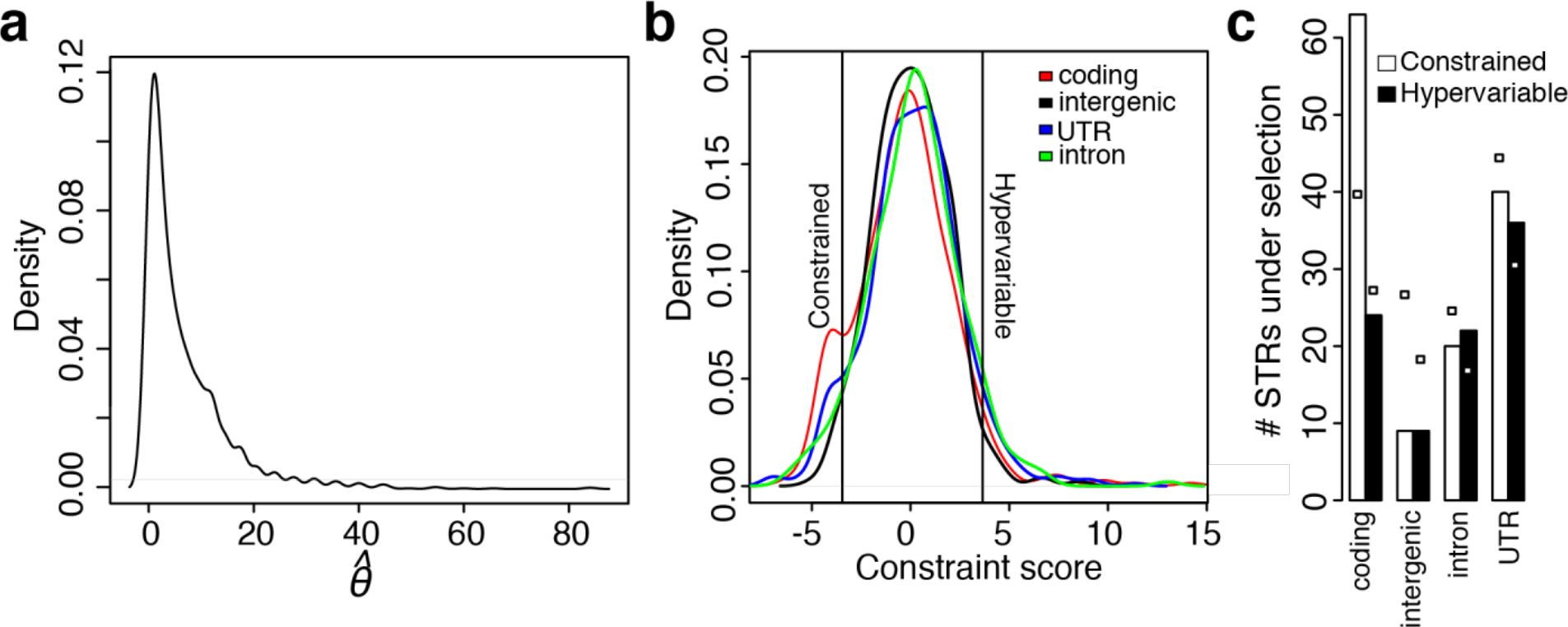
Detecting functionally constrained STRs. (a): The distribution of 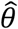 (estimated mutation rate (6)) across all genotyped STR loci. (b): Distribution of “selection scores” across all STRs, separated by locus category. Vertical lines indicate 2.5% and 97.5% quantiles of the distribution of intergenic STRs, which are used as thresholds for putative constraint and hypervariability respectively. (c): Constrained or hypervariable STRs separated by locus category. White boxes indicate the expected numbers for each bar, based on number of STRs in each locus category and number of STRs under different types of selection.

According to constraint scores, 132 STRs were less variable than expected under neutrality, suggesting purifying selection on these loci (Fig. 3c). Among these, coding STRs were overrepresented relative to their prevalence (OR = 2.4, p = 3.7*10^−6^, Fisher’s Exact Test; Figure 3c); this enrichment of coding STRs among constrained STRs agrees with our naïve analysis of invariant STRs above. Examples of constrained coding STRs included STRs encoding homologous polylysines adjoining the histone core in three different histone H2B proteins; notably, core histones are among the most conserved proteins across eukaryotes. Generally, coding STRs showing purifying selection encoded roughly half as many polyserines and twice as many acidic homopolymers as expected from the *A. thaliana* proteome (Supplementary Table 3) (37). While the interpretation of this association is unclear, it may be related to some structural role of such different classes of homopolymers in proteins. Although many more coding STRs are probably functionally constrained, our power to detect such constraints is limited by the size of the dataset. We also observed high conservation of some STRs in non-coding regions, though this is less interpretable given the ambiguous relationship between sequence conservation and regulatory function in *A. thaliana* (38). The most constrained intronic STR, in the *BIN4* gene, which is required for endoreduplication and normal development (39), shows a restricted allele frequency spectrum compared to similar STRs (Fig. 4a).

**Figure 4.**
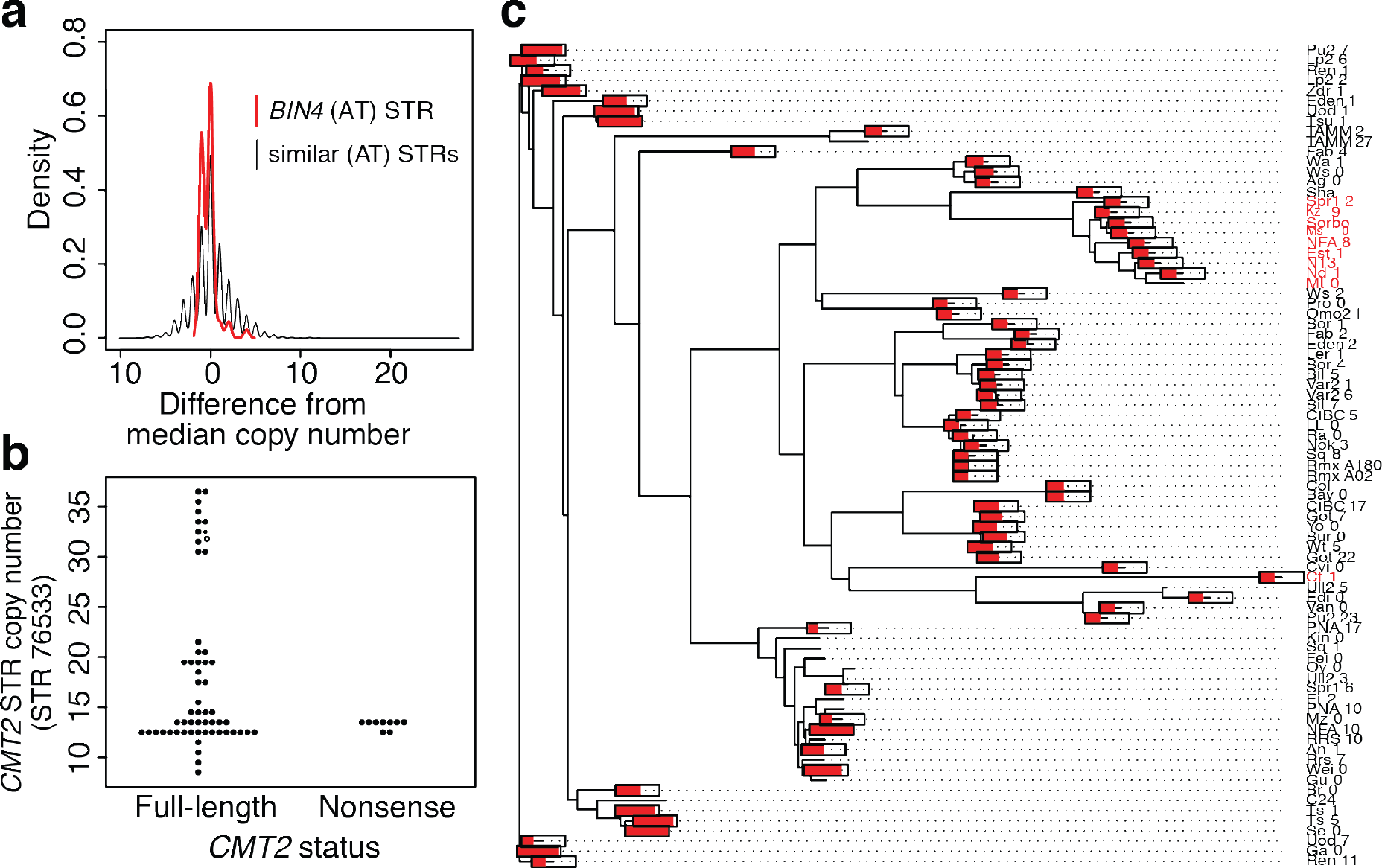
Non-coding STRs showing non-neutral variation. (a): *BIN4* intron STR is constrained relative to similar STRs. Allele frequency spectra are normalized by subtracting the median copy number (9 for the *BIN4* STR). All pure STRs with TA/AT motifs and a median copy number between 7 and 12 are included in the “similar STRs” distribution. (b): Lack of association between near-expansion *CMT2* STR alleles and previously described nonsense mutations. (c): Neighbor-joining tree of a 10KB region of *A. thaliana* chromosome 4 encompassing the *CMT2* gene across 81 strains with available data. Tip labels in red carry an adaptive nonsense mutation early in the first exon of *CMT2,* as noted previously (40). Red bars drawn on tips of the tree indicate the length of the *CMT2* intronic STR (as a proportion of its maximum length, 36.5 units). The bars are omitted for tips with missing STR data.

Hypervariable coding STRs (showing more alleles than expected) were too few for statistical arguments, but nonetheless showed several notable patterns (Supplementary Note, Supplementary Fig. 11). For example, 3/24 hypervariable coding STRs encoded polyserines in F-box proteins, suggesting that STR variation may serve as a mechanism of diversification in this protein family, which shows dramatically increased family size and sequence divergence in some plant lineages (41, 42). One non-coding hypervariable STR was associated with the *Chromomethylase 2* (*CMT2*) gene, which is under positive selection in *A. thaliana* (40). Specifically, CMT2 nonsense mutations in some populations are associated with temperature seasonality. We considered whether the extreme *CMT2* STR alleles might be associated with these nonsense mutations. Instead, these extreme alleles exclusively occurred in strains with full-length CMT2 (Fig. 4b). Strains with the common *CMT2* nonsense mutation form a tight clade in the *CMT2* sequence tree, whereas the *CMT2* STR length fluctuates rapidly throughout the tree and appears to converge on longer alleles independently in different clades (Fig. 4c). These convergent changes are consistent with a model in which the *CMT2* STR is a recurrent target of positive selection.

We further assessed whether STR conservation can be attributed to *cis*-regulatory function. The abundance of STRs in eukaryotic promoters (43), and their associations with gene expression (in *cis*) (19, 44), have suggested that STRs affect transcription, possibly by altering nucleosome positioning (44). We examined whether STRs near transcription start sites (TSSs) showed signatures of functional constraint (Fig. 5a). We found little evidence for reduced STR variation near TSSs, suggesting that *cis*-regulatory effects do not generally constrain STR variation in *A. thaliana.* Moreover, we found no relationship between constraint scores and location of STRs in accessible chromatin sites marking regulatory DNA (Fig. 5b).

**Figure 5.**
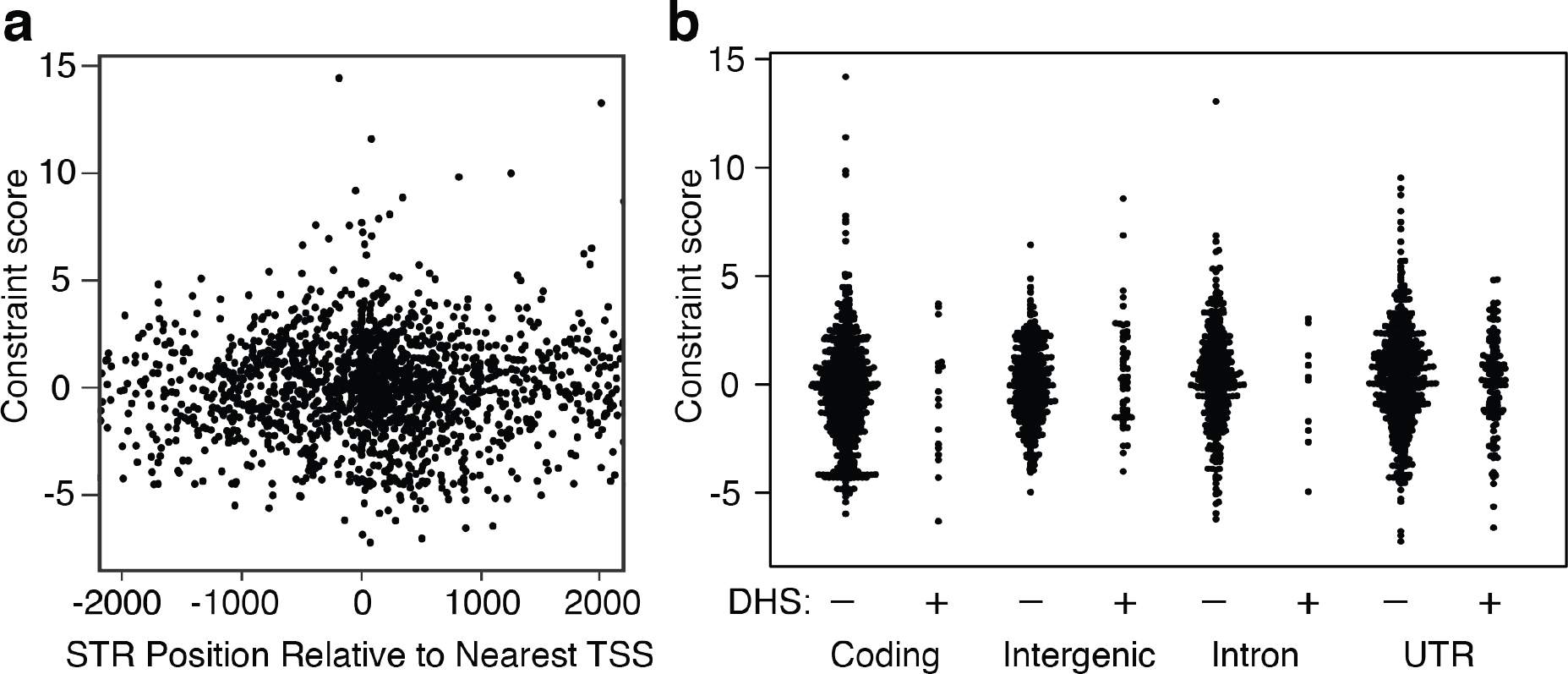
Relationship of STR constraint to gene regulatory elements. (a): Constraint score from Figure 3b plotted with respect to nearest TSS. (b): Constraint score from Figure 3b plotted with respect to STR annotations and presence of (putatively regulatory) DNase I hypersensitive sites (DHS).

### STRs yield numerous novel genotype-phenotype associations.

We next addressed the question of whether STR genotypes contribute new information for explaining phenotypic variation. It is often assumed that STR phenotype associations should be captured through linkage to common single nucleotide polymorphisms (SNPs) (45). This assumption is inconsistent with available data. Human STR data (45, 46) and simulation studies (47) indicate limited linkage disequilibrium (LD) between STRs and SNPs, making it unclear whether SNP markers can meaningfully tag STR genotypes. Indeed, we found little evidence of LD between STR and SNPs in *A. thaliana* (Fig. 6a). Moreover, the observed LD around STR loci declined with increasing STR allele number (Supplementary Fig. 12), consistent with an expected higher mutation rate at multiallelic loci (6, 47). This result suggests that STR-phenotype associations need to be directly tested rather than relying on linkage to
SNPs. *A. thaliana* offers extensive high-quality phenotype data for our inbred strains, which have been previously used for SNP-based association studies (48). We tested each polymorphic STR for associations across the 96 strains with each of 105 published phenotypes. For the subset of 32 strains for which RNA-seq data were available (49), we also tested for associations between STR genotypes and expression of genes within 1MB (*i.e.*, eQTLs; Supplementary Note). Although our power to detect eQTLs was limited by sample size, we detected 12 significant associations. The strongest association was between a STR residing in long noncoding RNA gene *AT4G07030* and expression of the nearby stress-responsive gene *AtCPL1* (Supplementary Fig 13, Supplementary Table 4).

We next focused on organismal phenotypes. Certain STRs showed associations with multiple phenotypes, and flowering time phenotypes were particularly correlated with one another (Fig. 6c,d). Similar to these patterns, SNPs have also shown associations with multiple phenotypes, and correlated flowering time phenotypes are among the strongest associations in the same strains (48). As in previous association studies using STRs (19), some inflation was apparent in test p-values compared to expectations, though the same tests using permuted STR genotypes showed negligible inflation (Fig. 6b, Supplementary Fig 14). Negligible inflation with permuted genotypes has been used previously to exclude confounding from population structure (19), which we will also presume here (Supplementary Note). We found 137 associations between 64 STRs and 25 phenotypes at stringent genome-wide significance levels (Methods; Supplementary Table 5). Given the low LD observed between STRs and other variants, STR variants may themselves be causal, rather than merely tagging nearby causal variants. Our analysis found plausible candidate genes, such as *COL9*, which acts in flowering time pathways and contains a flowering time-associated STR, and *RABA4B*, which acts in the salicylic acid defense response (50) and contains a STR associated with lesion formation.

**Figure 6.**
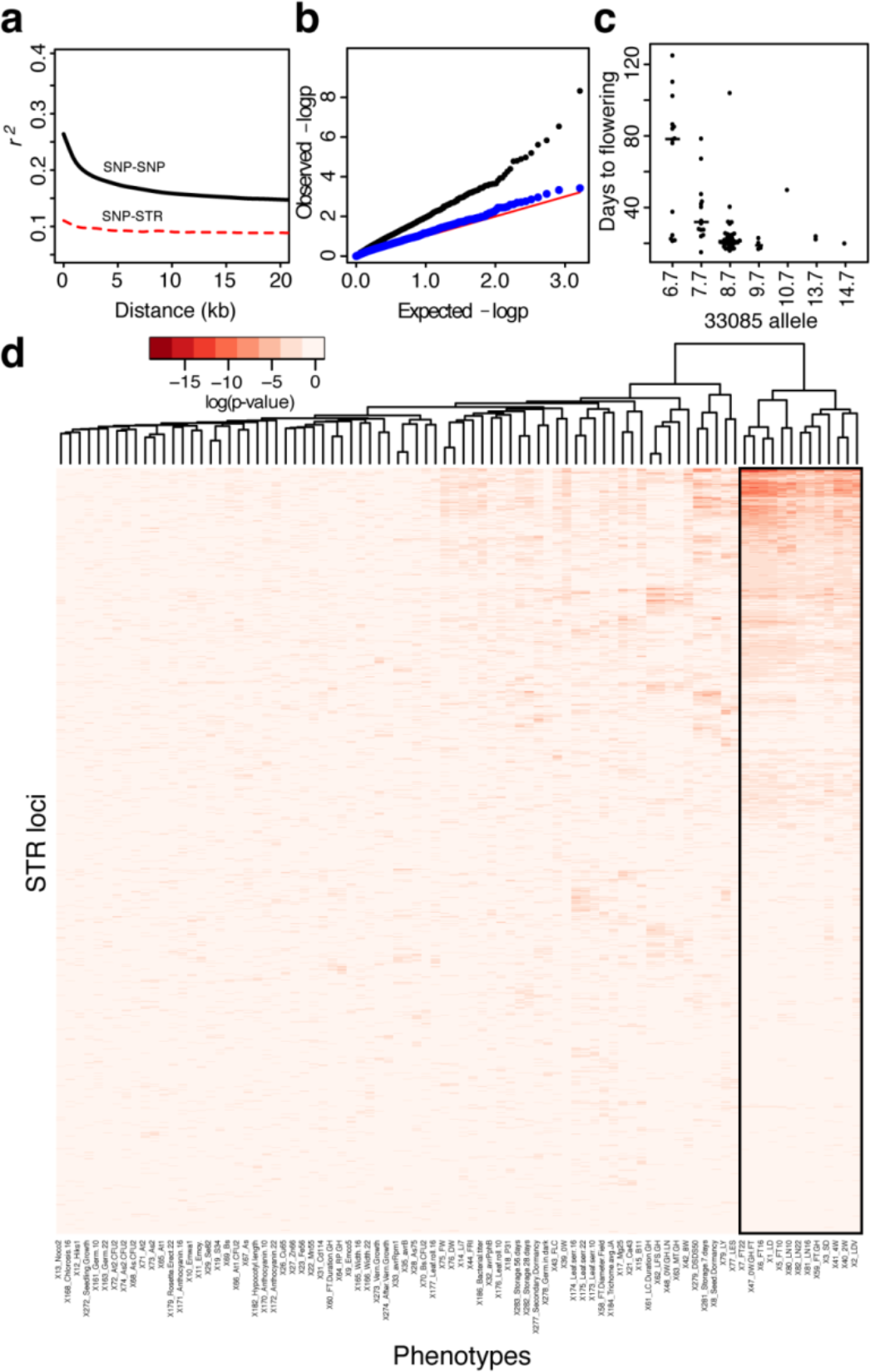
Diverse associations of STRs with quantitative phenotypes. (a): Multiallelic LD (51) estimates for STR and SNP loci. Lowess lines for each category are plotted. All values of r^2^ < 0.05 are omitted from lowess calculation for visualization purposes. (b): Quantile-quantile plot of p-values from tests of association between STRs and germination rate after 28 days of storage. (c): An example association between an STR (33085) and a phenotype (flowering time in long days after 4 weeks vernalization) in *A. thaliana* strains. Median of each distribution is indicated by a bar proportional in width to the number of observations. (d): Heatmap showing pairwise associations between STRs and phenotypes, summarized by the p-value from a linear mixed model, fitting STR allele as a fixed effect and kinship as a random effect. Both rows and columns are clustered, though the row dendrogram was omitted for clarity. STRs with genotype information in fewer than 25 strains are not displayed. Flowering time phenotypes are boxed in black.

We evaluated whether these associations might have been found using SNP-based analyses. We found that STR effects on phenotype are largely not accounted for by nearby SNP variation. Considering the strongest association for each STR, only 18 of the 64 STRs were near potentially confounding SNP variants, and most associations (14/18) were robust to adjustment for nearby SNP genotypes (48) (Supplementary Table 6, Supplementary Note). One notable exception was a STR closely linked to a well-known deletion of the *RPS5* gene in a hypervariable region of chromosome 1 (52) that causes resistance to bacterial infection (53). *RPS5* status is under balancing selection in *A. thaliana* (52). In this case, the association and the linkage are apparently strong enough (the STR is ~4KB upstream of the deletion) that this STR tags *RPS5’s* effect on infection. The observation of a STR tagging a hypervariable region leads us to speculate that STR variation holds information about genomic regions with complex mutational histories.

To assess the STR contribution to the variance of a specific trait, we performed a naïve variance decomposition of the long-day flowering phenotype into SNP and STR components, as represented by the loci showing associations with this trait. Our results suggested that STRs potentially contribute as much or more variance than SNPs to this phenotype (Supplementary Note, Supplementary Table 7). Estimated effect sizes for STR variants on this phenotype were similar to those of large-effect SNP variants (48) (Supplementary Fig 15).

Finally, we used mutant analysis to evaluate the two strongest flowering time associations, a coding STR in *AGL65* and an intronic STR in the uncharacterized gene *AT4G01390*; neither locus had been associated with flowering time phenotypes. We found that disruptions of both STR-associated genes conferred modest early flowering effects (by ~2 days and ~1 rosette leaf, p < 0.05 for each in linear mixed models; Supplementary Fig 16), supporting the robustness of our STR-phenotype associations. Taken together, our study suggests that STRs contribute substantially to phenotypic variation.

## Discussion

Our results imply that STRs contribute substantially to trait heritability in *A. thaliana*. There is little support for the hypothesis that STRs are “junk DNA”: STRs are apparently constrained by functional requirements, STR variation can disrupt gene function, and STR variation is associated with phenotypic variation. Considering that STR variation is represented poorly by nearby SNPs, the failure to directly ascertain STRs will mask most phenotypic consequences of such STR effects. If STR variation contributes to the genetic variance, as we argue here, the relevance of any genome-wide association study relying on SNP genotypes alone will be limited.

Compared to other classes of markers, STRs may exert an outsized influence on the phenotypic variance due to *de novo* mutations (V_m_; Table 1, (4)). Estimates of V_m_ from model organisms are on the order of 1%, but may vary substantially from trait to trait (54). STRs are good candidates for a substantial proportion of this variance, given their high mutation rate, residence in functional regions, and functional constraint demonstrated here (though we make no attempt to quantify V_m_ from STRs in this study). In previous work (55), we showed apparent copy number conservation of a STR in spite of a high mutation rate. In this case, deviation from the conserved copy number produced aberrant phenotypes. Our observation that constrained STRs are common suggests that STRs are a likely source of deleterious *de novo* mutations which are subsequently removed by selection.

The extent to which STRs affect phenotype is only partially captured in this study. Specifically, we assayed two STRs shown to cause phenotypic variation in prior transgenic *A. thaliana* studies (29, 56), but these STR loci did not show strong signatures of phenotypic association or of selection. This lack of ascertainment suggests that many more functionally important STRs exist in *A. thaliana* than we can detect with the analyses presented here. For example, the polyQ-encoding STR in the *ELF3* gene causes dramatic variation in developmental phenotypes (56), yet we find no statistical associations between this locus and phenotype across our 96 strains. In this case the lack of phenotype association is expected; *ELF3* STR alleles interact epistatically with several other loci (57), and thus would require increased power or more sophisticated analyses to detect associations. Indeed, we have argued that STRs are more likely than less mutable classes of genomic variation to exhibit epistasis (13). In consequence, we expect that the associations described in the present study are an underestimate of STR effects on phenotype. Moreover, our data are constrained by MIP technology, which limits the size and composition of STR alleles that we can ascertain (Figure 1a, Supplementary Figures 1-2).

Considering next a mechanistic perspective, the association we observe between intronic STR expansions and splice disruptions may be an important mechanism by which STRs contribute to phenotypic variation. In humans, unascertained diversity of splice forms contributes substantially to disease (58), and this diversity is larger than commonly appreciated (59). We demonstrate that this mechanism is common at least for expansions; future work should evaluate how tolerant introns are to different magnitudes of STR variation, as these effects on protein function may prove to be both large and relatively prevalent. The phenotypic contributions of loci with high mutation rate remain underappreciated, specifically in cases where such loci are difficult to ascertain with high-throughput sequencing. The results presented here argue that STRs are likely to play a substantial role in phenotypic variation and heritability. Accounting for the heterogeneity of different classes of genomic variation, and specifically variation in mutation rate, will advance our understanding of the genotype-phenotype map and the trajectory of molecular evolution.

## METHODS

### ONLINE METHODS

#### Probe design

We used TRF (60) (parameters: matching weight 2, mismatching penalty 5, indel penalty 5, match probability 0.8, indel probability 0.1, score ≥ 40 and maximum period 10) to identify STRs in the TAIR8 build of the *Arabidopsis thaliana* genome, identifying 7826 putative STR loci under 200bp (Supplementary File 2). We restricted further analysis to the 2409 loci with repeat purity >=89%. We chose 2307 STRs from among these, prioritizing STRs in coding regions, introns, or untranslated regions (UTRs), higher STR nit purity, and expected variability (VARscore (36)). We designed (61) molecular inversion probes (MIPs) targeting these STR loci in 180bp capture regions with 8bp degenerate tags in the common MIP backbone. For this purpose, we converted STR coordinates to the TAIR10 build and used the TAIR10 build as a reference genome. We used single nucleotide variants (SNVs) in 10 diverse *Arabidopsis thaliana* strains (62) to avoid polymorphic sites in designing MIP targeting arms. We filtered out MIPs predicted to behave poorly (MIPGEN logistic score < 0.7), discarded MIPs targeting duplicate regions, and substituted MIPs designed around SNVs as appropriate. We attempted to re-design filtered MIPs with 200bp capture regions using otherwise identical criteria. This yielded a final set of 2050 STR-targeting MIP probes (Supplementary File 3).

#### MIP and library preparation

These 2050 probes were ordered from Integrated DNA Technologies as desalted DNAs at the 0.2 picomole scale and resuspended in Tris-EDTA pH 8.0 (TE) to a concentration of 2 μM and stored at 4°C. We pooled and diluted probes to a final stock concentration of 1 nM. We phosphorylated probes as described previously (63). We performed DNA preparation from whole aerial tissue of adult *A. thaliana* plants. We prepared MIP libraries essentially as described previously(16, 63) using 100 ng *A. thaliana* genomic DNA for each of 96 *A. thaliana* strains.

#### Sequencing

We sequenced pooled capture libraries essentially as previously described(16) on NextSeq and MiSeq instruments collecting a 250bp forward read sequencing the ligation arm and captured target sequence, an 8bp index read for library demultiplexing, and a 50bp reverse read sequencing the extension arm and degenerate tag for single-molecule deconvolution. In each run, 10% of the sequenced library pool consisted of high-complexity whole-genome library to increase sequence complexity. For statistics and further details of data acquired for each library see Supplementary Tables 8 and 9.

#### STR annotation

We annotated STRs according to Araport11 (64), classifying all STRs as coding, intronic, intergenic, or UTR-localized, and indicating whether each STR overlapped with transposable element sequence. To identify regulatory DNA, we used the union of seven distinct DNaseI-seq experiments (65) covering pooled or isolated tissue types. For additional details, see the Supplementary Note.

#### Sequence analysis

Sequences were demultiplexed and output into FASTQ format using BCL2FASTQ v2.17 (Illumina, San Diego). We performed genotype calling essentially as described previously (16), with certain modifications (Supplementary Note, Supplementary Table 1).

Note that our *A. thaliana* strains are inbred, and more stringent filters and data processing would be necessary to account for heterozygosity. For information about comparison with the Bur-0 genome, see the Supplementary Note. Updated scripts implementing the MIPSTR analysis pipeline used in this study are available at https://osf.io/mv2at/?view_only=d51e180ac6324d2c92028b2bad1aef67.

#### Statistical analysis and data processing

We performed all statistical analysis and data exploration using R v3.2.1 (66). For plant experiments, we fit mixed-effects models using Gaussian (flowering) or binomial generalized linear models using experiment and position as random effects and genotype as a fixed effect.

#### STR expansion inference

We inferred STR expansions where the maximum copy number of an STR is at least three times larger than the median copy number of that STR. Various alleles of STR expansions were inspected manually in BAM files. Selected cases were dideoxy-sequenced and analyzed as described.

#### Plant material and growth conditions

Plants were grown on Sunshine soil #4 under long days (16h light : 8h dark) at 22°C under cool-white fluorescent light. T-DNA insertion mutants were obtained from ABRC (67) (Supplementary Table 11). For flowering time experiments, plants were grown in 36-pot or 72-pot flats; days to flowering (DTF) and rosette leaf number at flowering (RLN) were recorded when inflorescences were 1cm high. Results are combined across at least three experiments.

#### Gene expression and splicing analysis

We grew bulk seedlings of indicated strains on soil for 10 days, harvested at Zeitgeber time 12 (ZT12), froze samples immediately in liquid nitrogen, and stored samples at - 80°C until further processing. We extracted RNA from plant tissue using the SV RNA Isolation kit (including DNase step; Promega, Madison, WI), and subsequently treated it with a second DNAse treatment using the Turbo DNA-free kit (Ambion, Carlsbad, CA). We performed cDNA synthesis on ~500ng RNA for each sample with oligo-dT adaptors using the RevertAid kit (ThermoFisher, Carlsbad, CA). We performed PCR analysis of cDNA with indicated primers (Supplementary Table 10) and ~25 ng cDNA with the following protocol: denaturation at 95° 5 minutes, then 30 cycles of 95° 30 seconds, 55° 30 seconds, 72° 90 seconds, ending with a final extension step for 5 minutes at 72°. We gel-purified and sequenced electrophoretically distinguishable splice variants associated with STR expansions. Each RT-PCR experiment was performed at least twice with different biological replicates.

#### Population genetic analyses

For PCA, STRs with missing data across the 96 strains were omitted, leaving 987 STRs with allele calls for every strain. 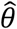 was estimated using the approximation 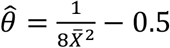, (6) where 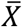 is the average frequency of all STR alleles at a locus. We computed multiallelic linkage disequilibrium estimates for SNP-SNP and SNP-STR locus pairs using MCLD (51). We downloaded array SNP data for the same lines (TAIR9 coordinates) from http://bergelson.uchicago.edu/wp-content/uploads/2015/04/call_method_75.tar.gz (68). For each locus, both SNP and STR, we computed linkage disequilibrium scores with 150 surrounding loci. To facilitate comparison, we computed lowess estimates of linkage only for those locus pairs in the plotted distance window in each case, and only for locus pairs with *r^2^* > 0.05.

#### Inference of conservation

STRs typed across 70 *A. thaliana* strains or fewer were dropped from this analysis, as the estimates of their variability were unlikely to be accurate, leaving 1825 STRs. We measured STR variation as the base-10 logarithm of the standard deviation of STR copy number (Supplementary Note). We used bootstrap aggregation (“bagging”) to describe a distribution of predictions as follows. An ensemble of 1000 support vector regression (SVR, fit using the *ksvm()* function in the kernlab package(69)) models was used to predict expected neutral variation of each STR as quantified by each measure (Supplementary Note). We used this distribution of bootstrapped predictions for intergenic STRs to compute putative conservation scores (Z-scores) for each STR. Scores below the 2.5% (Z < −3.46) and above the 97.5% (Z > 3.65) quantiles of intergenic STRs were considered to be putatively constrained and hypervariable respectively.

#### eQTL inference

We downloaded normalized transcriptome data for *A. thaliana* strains from NCBI GEO GSE80744 (49). We used the Matrix eQTL package (70) to detect associations, fitting also 10 principal components from SNP genotypes to correct for population structure. Following precedent (19), we fitted additive models assuming that STR effects on expression would be a function of STR copy number. We used Kruskal-Wallis rank-sum tests to test the null hypothesis of no association following correction.

#### Genotype-phenotype associations

We downloaded phenotype data from https://github.com/Gregor-Mendel-Institute/atpolydb/blob/master/miscellaneous_data/phenotype_published_raw.tsv. We followed precedent (48) in log-transforming certain phenotypes. In all analyses we treated STRs as factorial variables (to avoid linearity assumptions) in a linear mixed-effect model analysis to fit STR allele effects on phenotype as fixed effects while modeling the identity-by-state kinship matrix between strains (computed from SNP data) as a correlation structure for strain random effects on phenotype. We performed this modeling using the *Imekin()* function from the *coxme* R package (71). We repeated every analysis using permuted STR genotypes as a negative control to evaluate p-value inflation, and discarded traits showing such inflation. We used p < 10^−6^ as a genome-wide significance threshold commensurate to the size of the *A. thaliana* genome and the data at hand. For flowering time phenotypes we used a more stringent p < 10^−10^ threshold, as these phenotypes showed somewhat shifted p-value distributions (which were nonetheless inconsistent with inflation, according to negative controls). We identified potentially confounding SNP associations using the https://gwas.gmi.oeaw.ac.at/#/study/1/phenotypes resource, using the criterion that a SNP association must have a p <= 10^−4^ and be within roughly 100KB of the STR to be considered. We fit models including SNPs as fixed effects as before, and performed model selection using AICc (72). Additional details about association analyses are in the Supplementary Note.

#### Data and code availability

MIP sequencing data are available in FASTQ format at the Sequence Read Archive, under project number PRJNA388228. Analysis scripts are provided at https://osf.io/5jm2c/?view_only=324129c85b3448a8bd6086263345c7b0, along with data sufficient to reproduce analyses.

## ACKNOWLEDGMENTS

We thank Keisha Carlson, Alberto Rivera, and members of the Queitsch lab for technical assistance and important conversations. We thank Evan Boyle, Choli Lee, Matthew Snyder, and Jay Shendure for assistance and advice concerning MIP design and use. We thank the Dunham Lab and the Fields Lab for access to and help with sequencing instruments. We thank UW Genome Sciences Information Technology for high-performance computing resources. This work was supported in part by NIH New Innovator Award DP2OD008371 to CQ.

